# Proteomics Analysis of Differential Expression of Lung Proteins in Response to Highly Pathogenic Avian Influenza Virus Infection in Chicken

**DOI:** 10.1101/2019.12.16.877456

**Authors:** Periyasamy Vijayakumar, Ashwin Ashok Raut, Santhalembi Chingtham, Harshad V Murugkar, Diwakar D. Kulkarni, Vijendra Pal Singh, Anamika Mishra

**Affiliations:** Pathogenomics Laboratory, ICAR-National Institute of High Security Animal Diseases, OIE Reference lab for Avian Influenza, Bhopal - 462021, Madhya Pradesh, India; ICAR -National Institute of High Security Animal Diseases, OIE Reference lab for Avian Influenza, Bhopal - 462021, Madhya Pradesh, India

**Keywords:** Avian influenza virus, proteome, molecular pathways, proteomic determinants, chicken, lung tissues

## Abstract

Elucidation of molecular pathogenesis underlying virus-host interaction is important for the development of new diagnostic and therapeutic strategies against highly pathogenic avian influenza (HPAI) infection in chicken. However, chicken HPAI viral pathogenesis is not completely understood. To elucidate the intracellular signaling pathways and critical host proteins associated with influenza pathogenesis, we characterized the lung proteome of chicken infected with HPAI H5N1 virus (A/duck/India/02CA10/2011/Agartala). The chicken mass spectra data sets comprised1, 47, 451 MS scans and 19, 917 MS/MS scans. At local FDR 5% level, we identified total 3313 chicken proteins with presence of at least one unique peptide. At 12 hrs, 247 proteins are downregulated while 1754 proteins are downregulated at 48 hrs indicating that the host has succumbed to infection. There is expression of proteins of the predominant signaling pathways, such as TLR, RLR, NLR and JAK-STAT signaling. Activation of these pathways is associated with cytokine storm effect and thus may be the cause of severity of HPAI H5N1 infection in chicken. Further we identified proteins like MyD88, IKBKB, IRAK4, RELA, and MAVS involved in the critical signaling pathways and some other novel proteins(HNF4A, ELAVL1, FN1, COPS5, CUL1, BRCA1 and FYN) as main hub proteins that might play important roles in influenza pathogenesis in chicken. Taken together, we characterized the signaling pathways and the proteomic determinants responsible for disease pathogenesis in chicken infected with HPAI H5N1 virus.

## Introduction

The control measures to any emerging and re-emerging infection, requires an adequate understanding of virus-host interaction (Fauci, 2006). The complete understanding of virus-host interaction will provide important clues for the development of new diagnostic and therapeutic strategies against emerging diseases. The high-throughput functional genomics approach provides a deeper understanding of emerging disease by encompassing both the pathogen and the host response. Virus-host interactions are multidimensional in nature, and this interaction alters various host components such as transcriptome, proteome, miRNA, metabolome, and lipidome (Mishra et al., 2017). In past years, most of the high-throughput studies on H5N1-host interactions have focused at the transcriptome level.

However, transcriptomic studies do not reveal any posttranscriptional regulation, posttranslational modifications, and protein–protein interactions (Josset et al., 2013). The virus-host interaction involves the alternation of hundreds to thousands of host proteins. Hence, applying high throughput proteomics approach for the analysis of virus-induced innate immune responses in conjunction with the transcriptomic analyses promises better understanding of molecular mechanism of influenza pathogenesis (Zak et al., 2014). Proteomics refers to the large-scale study of protein expression, protein-protein interactions or posttranslational modifications based on high-resolution mass spectrometry (Gingras et al., 2007; Altelaar et al., 2013).

Proteomics studies of influenza virus infection in macaques (Brown et al., 2010), mice (Kumar et al., 2014), continuous cell lines (Vester et al., 2009; Coombs et al., 2010; Kummer et al., 2014), chicken (Zou et al., 2010; Li et al., 2017), dogs (Su et al., 2015) and primary human cells (Liu et al., 2008; Lietzen et al., 2011; Kroeker et al., 2012; Liu et al., 2012) have provided information on alteration of host proteome at cellular and organism levels. Recent studies of chicken proteome response against H5N1 infection revealed several alterations of cytoskeleton, metabolic process, cellular component, and transcription regulation proteins (Zou et al., 2010; Li et al., 2017). However, detailed knowledge of signalling pathways and proteomic determinants responsible for the HPAI H5N1 viral pathogenesis in avian species is not known. In this study, the intracellular signaling pathways and critical host proteins responsible for influenza pathogenesis in chicken lungs infected with HPAI H5N1 virus are characterized at the proteome level.

## Materials and Methods

### Experimental infection of chicken

Six weeks old healthy domestic chicken, sero-negative for Avian Influenza virus (AIV) were used for this study. The animal experiments were approved by the Institutional Animal Ethics Committee of ICAR-NIHSAD (Approval no. 68/IAEC/HSADL/12 dated 11.05.2012) and all the experiments were conducted in biosafety level 3 containment facility of ICAR-National Institute of High Security Animal Diseases, Bhopal, India. The chicken were separated into four groups (n=5 birds/group). Among the four groups, three groups were intranasally inoculated with 10^6^ EID_50_ of H5N1 virus (A/duck/India/02CA10/2011/Agartala) and one group (control) was inoculated with PBS. The birds were observed daily for clinical signs. Lung tissues were collected from five birds from each infected group at 12hr, 24 hr, and 48hr post-infection. Lung tissues were collected from the control group at 12hr post-inoculation. The tissues were snap chilled in Liquid Nitrogen and stored at −80°C until protein extraction. Avian influenza virus infection of lung tissues was confirmed by virus isolation upon inoculation in embryonated chicken eggs (ECEs) and RT-PCR.

### Protein extraction

150mg lung tissue from each sample was washed in 50mM NH_4_HCO_3_ washing buffer. The lung tissue was cut into small pieces and 650 μl of SDS protein extraction lysis buffer [0.1% SDS (Invitrogen); 50mM NH_4_HCO_3_ (Sigma); 1X cOmplete™ Protease Inhibitor Cocktail (Roche-11836145001)] was added. Tissue samples were homogenized in LZ-Lyser homogenizer at 30 HZ for 2 min. After complete homogenization, the tissue lysate was incubated on ice for 90 mins for complete protein lysis. The lysate was centrifuged at 20,000g for 60 mins at 4°C and the supernatant was collected. The supernatants were immediately snap heat treated at 56°C for 30 min in a dry bath for inactivation of HPAIV H5N1 in the protein extracts. All the heat treated samples were stored at −80°C for mass spectrometry analysis.

### Sample preparation for LC-MS analysis

Lung protein quality was checked by 8% SDS-PAGE. A pool for each time point was prepared by pooling 50μg protein each from 3 best samples at that time point. Protein samples were reduced for 1 hr at 95°C in 100mM dithiothreitol solution followed by alkylation for 45min by 55mM iodacetamide in the dark at RT. Trypsin was added to all protein samples at a 1:20 (wt/wt) trypsin-to-protein ratio for overnight at 37°C. After trypsin digestion, the sample quality was again checked by SDS PAGE. Digested peptide samples were concentrated to 50µl total volume using a centrifugal vacuum. The non-infected and infected peptide samples were labeled with iTRAQ 4-Plex (P/N: 4352135) reagents. The labels used for sample pools of different groups were as follows: iTRAQ label 114-control; iTRAQ label 115-12hr; iTRAQ label 116-24hr; iTRAQ label 117-48hr. These iTRAQ labeled samples were pooled and then purified using a strong cation exchange (SCX) chromatography. The fractions from SCX chromatography were collected and pooled according to RT values. The pooled fractions were vacuum dried and dissolved in 10μL of 0.1% formic acid. The sample was injected (1μL) onto C18 Nano-LC column for separation of peptides followed by mass spectrometry analysis on the WATERS Q-TOF instrument.

### Bioinformatics analysis

The WATRES specific raw data set files were converted to proteomics standard format mzXML format by proteowizard tool MSconvert with default parameters option (Chambers et al., 2012). The database search and other downstream bioinformatic analysis were done in Trans-Proteomic Pipeline (TPP) (Deutsch et al., 2010). TPP includes modules for validation of database search results, quantitation of isotopically labeled samples, and validation of protein identifications. MS/MS ion spectra were searched against chicken UniProt reference database using comet search engine (Eng et al., 2013). All 17719 proteins that comprise the chicken UniProt reference database and 11 protein sequences of HPAIV H5N1 virus (A/duck/India/02CA10/2011/Agartala from NCBI protein database (http://www.ncbi.nlm.nih.gov/protein/?term=txid1004759[Organism:noexp) were downloaded. The following parameters were used for database search, precursor/peptide mass tolerance-1.8Da, fragment tolerance −1.6Da, fixed modification – carboxamidomethylation for cys (C) and iTRAQ N terminal, Variable modification-oxidation (M) and phosphorylation of (STY), and number of missed cleavage 2. The Peptide-Spectrum assignments of comet search engine were validated with PeptideProphet and iProphet tool of TPP (Keller et al., 2002). Protein identifications were validated with ProteinProphet tool of TPP based on PeptideProphet or iProphet results (Nesvizhskii and Aebersold, 2004). Proteins identified from ProteinProphet result were filtered based on protein probability above 0.95 (local FDR 5% level) and contained at least 1 unique peptide. This filtered protein list was used for further downstream functional analysis. Up and downregulation of particular protein was calculated as infected sample intensity divided by control sample intensity (i.e. 115-12hr/114-control).Likewise for all proteins and specific post-infection time interval, fold change values were calculated. Functional classification of the proteins was performed for gene ontology (GO) in Database for Annotation, Visualization and Integrated Discovery (DAVID) (Huang et al., 2009) and pathway analysis in Kyoto Encyclopedia of Genes and Genomes (KEGG) (www.genome.jp/kegg/) (Kanehisa et al., 2017). Heatmap generated using ‘Clustvis’ web tool (Metsalu et al., 2015). We used online web server NetworkAnalyst for construction of protein-protein interaction (PPI) networks (Xia et al., 2014). The main driving or hub proteins were identified on the basis of two topological measures, degree centrality and betweenness centrality.

### Meta-analysis of transcriptome datasets

For meta-analysis of chicken transcriptome datasets, we selected the microarray datasets of 2 independent studies. Hu et al. (2015) studied immune response of primary chicken lung cells infected with two HPAI H5N1 viruses using microarray technology (Hu et al., 2015). Ranaware et al. (2016) studied global immune response of chicken infected with HPAI H5N1 (A/duck/India/02CA10/2011) virus infection (Ranaware et al., 2016). The microarray datasets were analysed in GEO2R online tool of the NCBI (http://www.ncbi.nlm.nih.gov/geo/geo2r/).The original submitter-supplied processed microarray data tables were identified using the GEO query. Then, we identified control and test group samples and then samples belonging to each group were assigned. Further, the logFC, p-value and adj. p-value were calculated by limma R package. Adjusted p-values below 0.05 were used as threshold to find differentially expressed genes.

Bayesian networks were reconstructed for TLR, RLR, IL1R, NLR and JAK-STAT signaling pathways using ‘bnlearn’ package (Scutari, 2017). The networks were constructed based on intensity values of microarray dataset of our pervious published work (Ranaware et al., 2016). Bayesian network structure was learned from transcriptomic dataset with prior knowledge using Hill Climbing (HC) algorithm. HC is a score-based structure learning algorithm. After learning the network structure, the conditional probability tables (CPTs) at each node were found by running bn.fit function. Bn.fit runs the EM algorithm to learn CPT for different nodes in the learned graph.

## Results and discussion

Virus-host interactions are multidimensional, they include the alterations in transcriptome, proteome, metabolome and lipidome of the host. In recent years, most of the high-throughput studies have focused majorly on the host transcriptomic responses and other host components have received less attention. This study presents a comprehensive characterization of the lung proteome, critical signaling pathways and proteomic determinants responsible for disease pathogenesis at the proteome level in chicken lung tissues infected with HPAI H5N1 virus over different time points post-infection.

### Clinical signs

Chicken in the control group did not show any clinical signs during the experimental period. In the test group, the birds were normal up to 12 hr post-infection. Mild clinical symptoms like depression, decreased feed as well as water consumption and ruffled feathers were observed at 24 hr post-infection. The clinical symptoms like dullness, lacrimation, cyanotic combs and wattles, edema and red discoloration of the shanks and feet were seen in the birds at 48 hr post-infection.

### Raw mass spectra dataset analysis

Total 15 fractions of WATERS QTOP raw data sets were generated, with data size 48 GB, including 1, 47,451 MS scans and 19,917 MS/MS scans. Total 19,917 MS/MS spectra were searched against chicken protein database by comet search engine. The iProphet algorithm identified 17,273 unique peptides and 8,516 unique proteins at minimum probability threshold of above 0.05 and minimum above 7 amino acids. In order to increase the accuracy of validation of peptides and proteins, we applied local FDR 5% level (probability cut off above 0.95) in the ProteinProphet output. At higher probability threshold cut off, we identified total 3313 proteins with the presence of at least one unique peptide. ProteinProphet predicted sensitivity and error rate information is shown in Figure S1. This is the highest protein identification reported till date in chicken. Our proteomic approach identified H5N1 viral peptides such as NA, NP and PB1 in the chicken lung proteome. NA is a sialidase responsible for releasing sialic acid from glycoprotein and glycolipid sialoconjugates of bound influenza virus to assist virus release (Wagner et al., 2001). NP is an important viral protein responsible for the packaging of the viral RNA and also shown to be involved in many aspects of influenza viral replication (Portela and Digard, 2002; Wasilenko et al., 2008). PB1 viral protein has been shown to be associated with the high pathogenicity of H5N1 viruses infection in ducks (Hulse-Post et al., 2007). Influenza viral infection status of lung tissues was confirmed by virus isolation in embryonated egg inoculation and RT-PCR method as well as by identification of viral peptides in lung proteome dataset.

### Differential protein expression analysis

At 12 hr post-infection, 820 proteins were upregulated and 2493 proteins were downregulated in chicken lung tissues. A total of 827 proteins were upregulated and 2441 proteins were downregulated at 24 hr post-infection. At 48 hr post-infection, 693 proteins were upregulated and 2620 proteins were downregulated in chicken lung proteome (Table 1). A total of 470 and 2235 proteins were found to be commonly upregulated and downregulated in all time intervals post-infection in chicken lungs infected with H5N1 virus, respectively (Figure 1). The protein profile showed that 70, 70 and 101 proteins were exclusively up regulated in chicken lung tissue at 12hr, 24hr and 48hr post-infection, respectively (Figure 1). Downregulation of 87, 35 and 245 proteins were observed exclusively at 12 hr, 24hr and 48hr post-infection, respectively (Figure 1). The fold change value of the upregulated proteins ranged from 42 to 1. Interestingly, at 48 hr time point (fold change value below1.5), higher number of proteins were downregulated (n=1754) as compare to 12 hr time point (n=247) post-infection condition (Table 1, Figure 2). This result indicates that most of the host proteins were downregulated at the later stage of infection.

**Table 1.** Differential protein expression analysis in chicken infected with HPAI H5N1 virus.

| Condition | Upregulated proteins | Downregulated proteins | Upregulated proteins (>1.5 fold) | Downregulated Proteins (<1.5 fold) |
| --- | --- | --- | --- | --- |
| 12hr interval | 820 | 2493 | 138 | 247 |
| 24hr interval | 872 | 2441 | 157 | 222 |
| 48hr interval | 693 | 2620 | 173 | 1754 |

**Fig 1.**
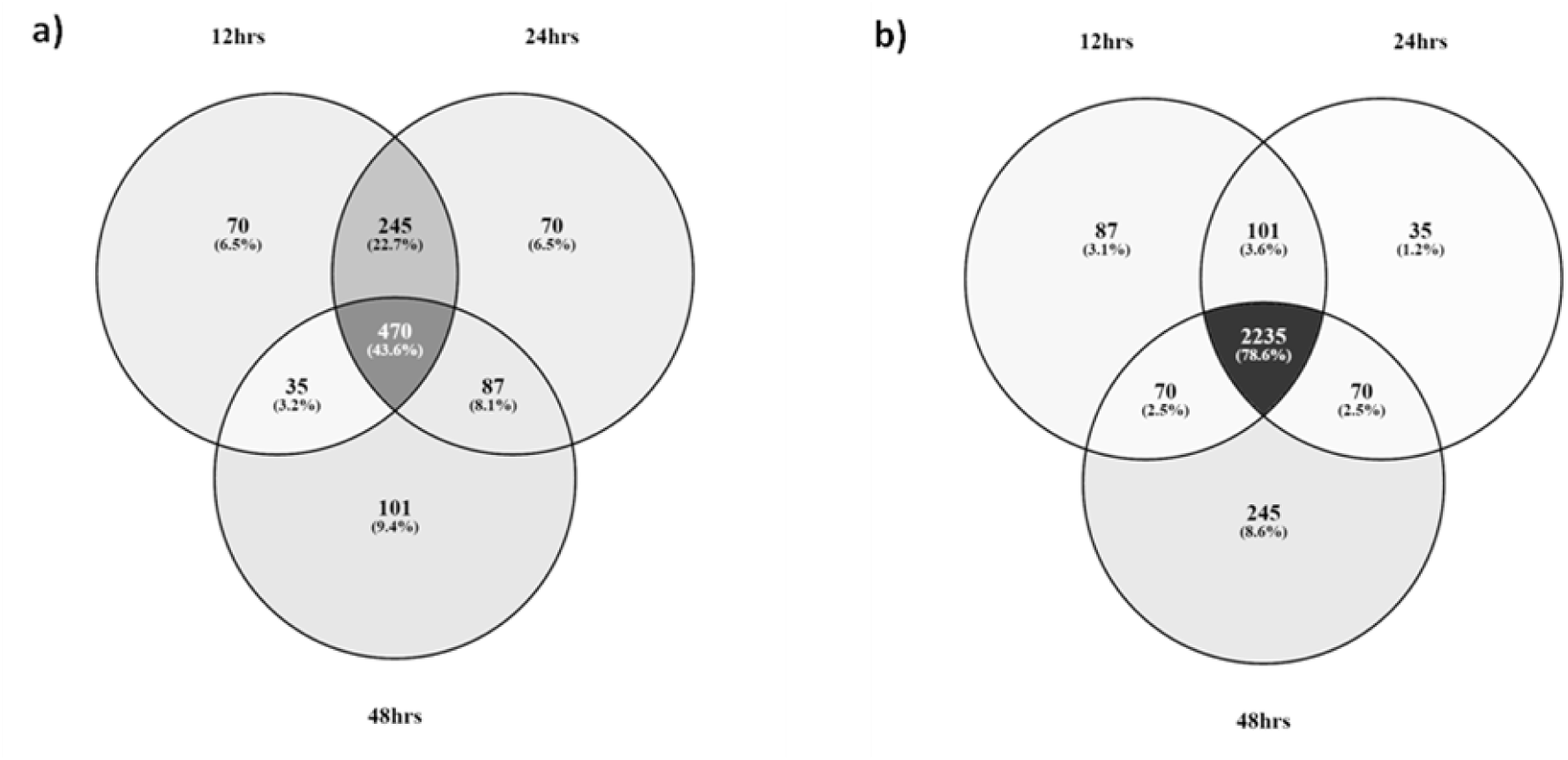
Comparative analysis of upregulated (a) and downregulated (b) proteins expression changes at different time intervals in chicken lung tissues.

**Fig 2.**
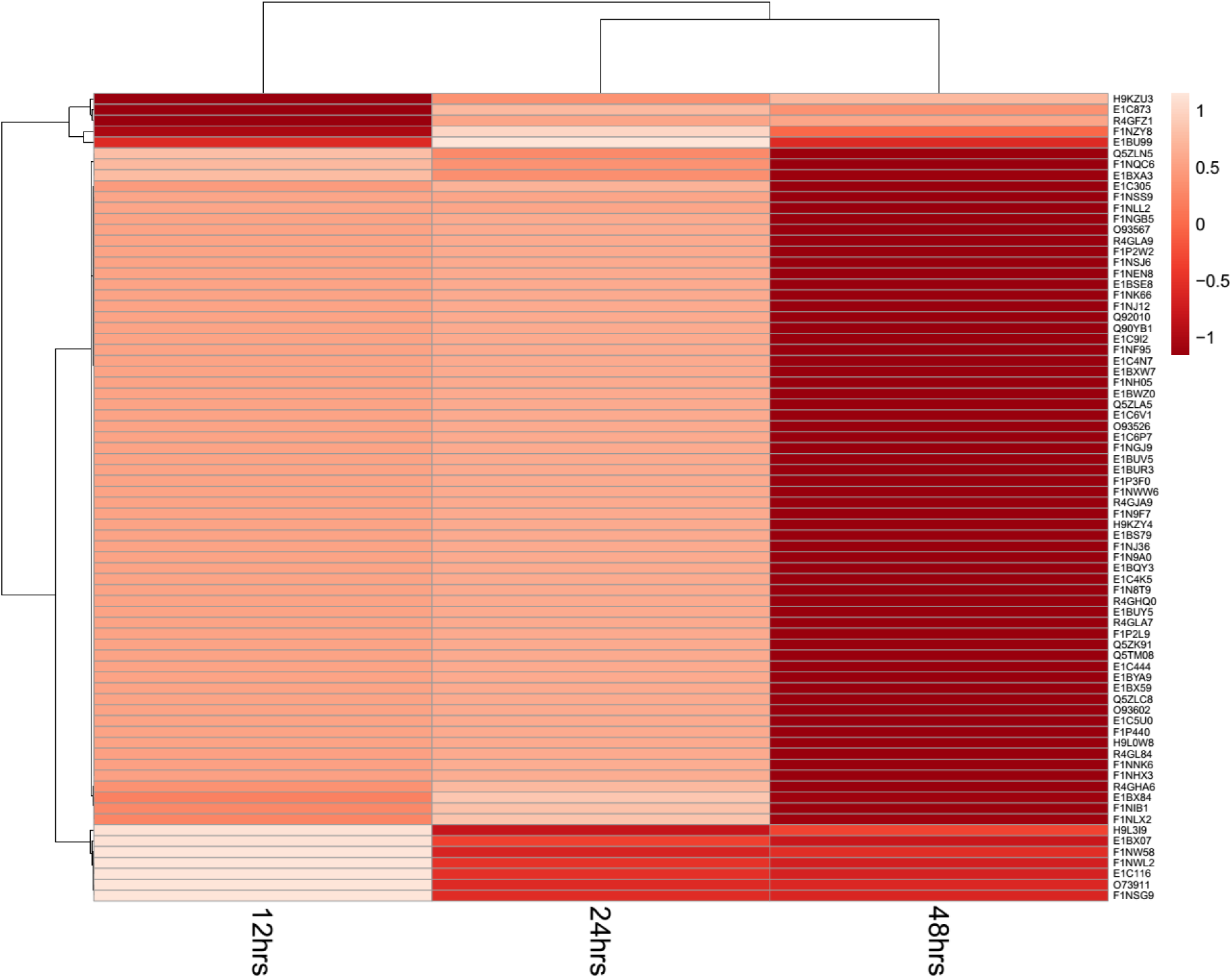
Heatmap of differentially expressed proteins of HPAIV infected lung tissues. The expression levels are visualized using a gradient colour scheme.

Gene ontology analysis of the commonly upregulated and downregulated lung tissue proteins of H5N1 infected chicken lung enriched following GO terms such as cytoskeleton, regulation of cell cycle, regulation of protein kinase activity etc. (Table 2). The upregulation (KRT6A, MCPH1, MYH7, MICAL1 and MICAL3) and downregulation (ACTL9, CTNNB1, DNM1, FILIP1L, and MYO1B) of cytoskeletal proteins were observed in the lung tissue of chicken. Cytoskeletal proteins have been reported to interact with viral proteins to regulate viral replication and assembly as well as the transport of viral components in the cell (Avalos et al., 1997; Radtke et al., 2006). Similar associations of cytoskeletal proteins with influenza infection condition are also reported previously (Coombs et al., 2010; Sui et al., 2014; Su et al., 2015). The cyclin-dependent kinases (CDKs) such as CDK13, DDB1, DRD3, FOXG1, and TCF3 that are involved in the regulation of cell cycle were all upregulated in chicken lung proteome. Söderholm *et al*. (2016) reported that the cyclin-dependent kinase activity is required for efficient viral replication and for activation of the host antiviral responses (Soderholm et al., 2016). Protein kinase activity associated proteins, namely, JAK3, MARK3, TBK1, EIF2AK3, PRKCA, and TRPM6were downregulated in the chicken lung tissues. Differential expression of proteins including signal transduction molecules, kinases and other biochemical metabolism related enzymes that are associated with the repair of damaged lung tissues are reported in dogs infected with influenza virus infection (Su et al., 2015). In addition, some apoptosis and tumor-associated proteins (BCL6, FAF1, TBX5, AKAP13, TNFSF10 and TGFB1) were also identified in the chicken lung proteome. Further we did GO term analysis of proteins that were exclusively expressed in particular post-infection condition in chicken (Table 3). This analysis result provides information how the disease progress from onset of disease to outcome of disease in chicken. At 12 hr interval, most of cellular homeostasis were affected and at 48 hr post-infection condition the host activated critical pathways such as Influenza A, chemokine signaling pathway, Jak-STAT signaling pathway, apoptosis, MAPK signaling pathway (Table 3). These results indicate that the virus initially inhibit the cellular homeostasis and activates the critical pathways at the later stage of infection.

**Table 2.**
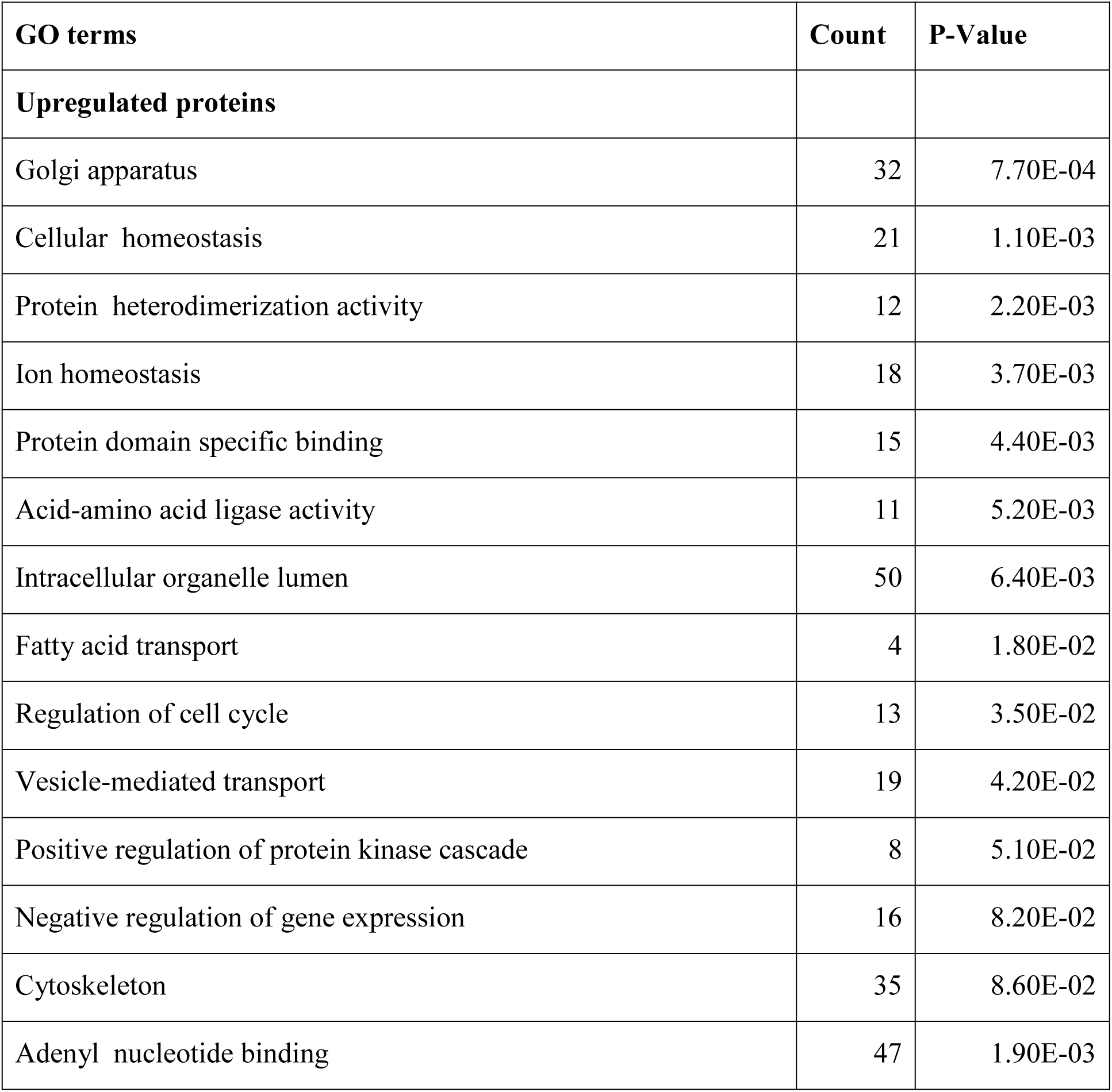
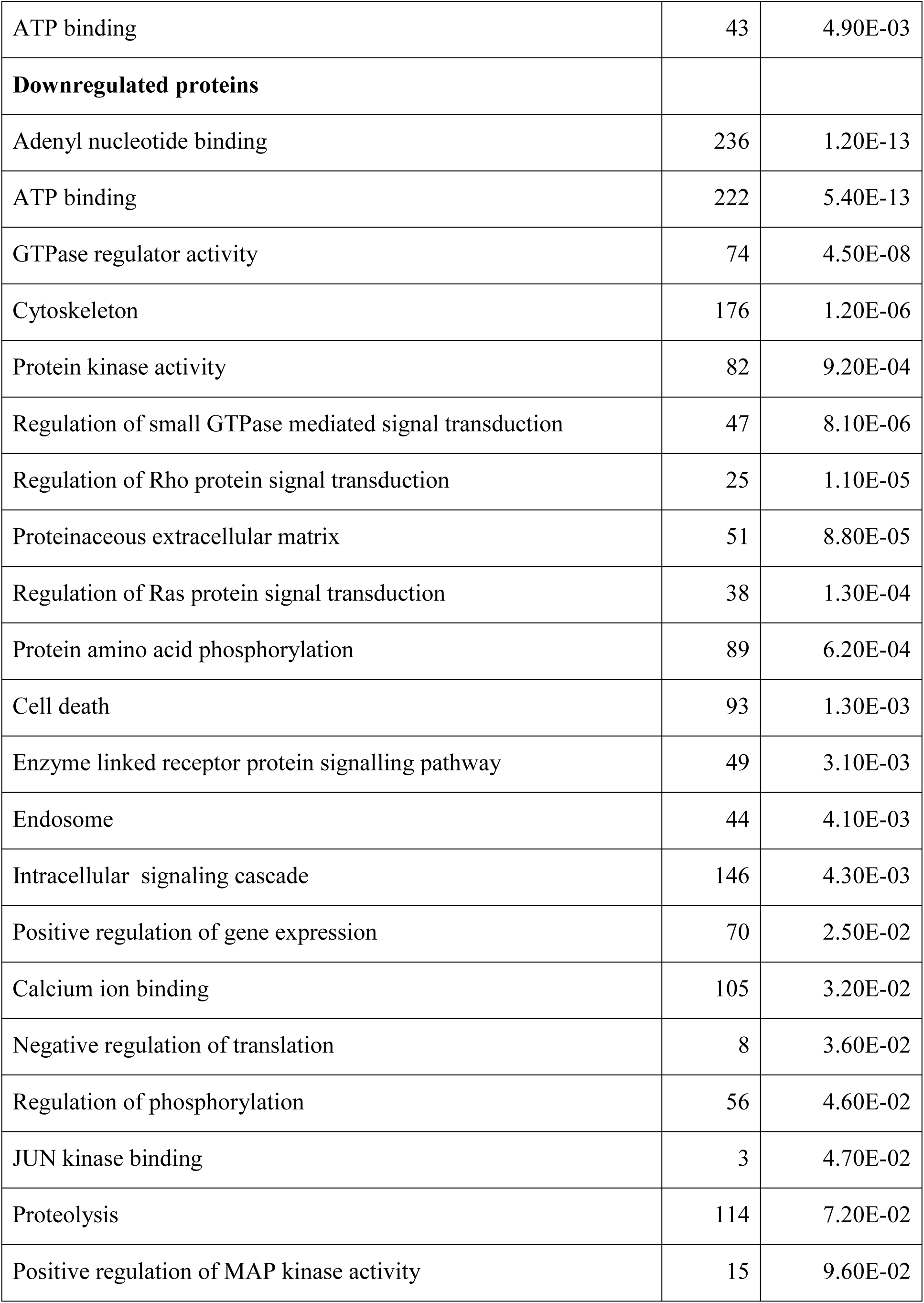
Gene Ontology term analysis of commonly upregulated and downregulated lung tissue proteins in chicken infected with HPAI H5N1 virus.

| GO terms | Count | P-Value |
| --- | --- | --- |
| **Upregulated proteins** |  |  |
| Golgi apparatus | 32 | 7.70E-04 |
| Cellular homeostasis | 21 | 1.10E-03 |
| Protein heterodimerization activity | 12 | 2.20E-03 |
| Ion homeostasis | 18 | 3.70E-03 |
| Protein domain specific binding | 15 | 4.40E-03 |
| Acid-amino acid ligase activity | 11 | 5.20E-03 |
| Intracellular organelle lumen | 50 | 6.40E-03 |
| Fatty acid transport | 4 | 1.80E-02 |
| Regulation of cell cycle | 13 | 3.50E-02 |
| Vesicle-mediated transport | 19 | 4.20E-02 |
| Positive regulation of protein kinase cascade | 8 | 5.10E-02 |
| Negative regulation of gene expression | 16 | 8.20E-02 |
| Cytoskeleton | 35 | 8.60E-02 |
| Adenyl nucleotide binding | 47 | 1.90E-03 |
| ATP binding | 43 | 4.90E-03 |
| **Downregulated proteins** |  |  |
| Adenyl nucleotide binding | 236 | 1.20E-13 |
| ATP binding | 222 | 5.40E-13 |
| GTPase regulator activity | 74 | 4.50E-08 |
| Cytoskeleton | 176 | 1.20E-06 |
| Protein kinase activity | 82 | 9.20E-04 |
| Regulation of small GTPase mediated signal transduction | 47 | 8.10E-06 |
| Regulation of Rho protein signal transduction | 25 | 1.10E-05 |
| Proteinaceous extracellular matrix | 51 | 8.80E-05 |
| Regulation of Ras protein signal transduction | 38 | 1.30E-04 |
| Protein amino acid phosphorylation | 89 | 6.20E-04 |
| Cell death | 93 | 1.30E-03 |
| Enzyme linked receptor protein signalling pathway | 49 | 3.10E-03 |
| Endosome | 44 | 4.10E-03 |
| Intracellular signaling cascade | 146 | 4.30E-03 |
| Positive regulation of gene expression | 70 | 2.50E-02 |
| Calcium ion binding | 105 | 3.20E-02 |
| Negative regulation of translation | 8 | 3.60E-02 |
| Regulation of phosphorylation | 56 | 4.60E-02 |
| JUN kinase binding | 3 | 4.70E-02 |
| Proteolysis | 114 | 7.20E-02 |
| Positive regulation of MAP kinase activity | 15 | 9.60E-02 |

**Table 3.** Functional annotation analysis of proteins exclusively expressed in chicken at different time points post infection with HPAI H5N1 virus.

| Pathway activation | No. of proteins | P Value |
| --- | --- | --- |
| **12 hr post infection** |  |  |
| Cell cycle | 31 | 2.88E-26 |
| mRNA surveillance pathway | 21 | 4.95E-18 |
| RNA transport | 24 | 2.62E-17 |
| p53 signaling pathway | 11 | 1.22E-07 |
| Pathways in cancer | 13 | 0.0128 |
| **24 hr post infection** |  |  |
| Cell cycle | 12 | 1.17E-08 |
| Gap junction | 8 | 7.31E-06 |
| RNA degradation | 6 | 6.13E-05 |
| T cell receptor signaling pathway | 7 | 0.000127 |
| Fc epsilon RI signaling pathway | 6 | 0.000215 |
| ErbB signaling pathway | 6 | 0.000485 |
| Phagosome | 4 | 0.00326 |
| Fc gamma R-mediated phagocytosis | 5 | 0.00529 |
| Chemokine signaling pathway | 7 | 0.00615 |
| B cell receptor signaling pathway | 4 | 0.0112 |
| Neurotrophin signaling pathway | 5 | 0.0141 |
| Natural killer cell mediated cytotoxicity | 5 | 0.0221 |
| Focal adhesion | 6 | 0.0287 |
| **48 hr post infection** |  |  |
| RNA transport | 47 | 1.77E-18 |
| T cell receptor signaling pathway | 37 | 6.08E-15 |
| Regulation of actin cytoskeleton | 52 | 1.15E-14 |
| Focal adhesion | 54 | 4.67E-14 |
| Pathways in cancer | 70 | 1.13E-13 |
| ErbB signaling pathway | 31 | 6.41E-12 |
| Adipocytokine signaling pathway | 26 | 6.92E-12 |
| B cell receptor signaling pathway | 28 | 1.91E-11 |
| Neurotrophin signaling pathway | 35 | 3.79E-10 |
| mRNA surveillance pathway | 26 | 6.14E-09 |
| Chemokine signaling pathway | 41 | 7.57E-08 |
| Influenza A | 28 | 1.68E-07 |
| Cell cycle | 30 | 3.95E-07 |
| Jak-STAT signaling pathway | 25 | 1.62E-06 |
| Fc epsilon RI signaling pathway | 21 | 1.86E-06 |
| mTOR signaling pathway | 14 | 2.72E-05 |
| Toll-like receptor signaling pathway | 20 | 0.000377 |
| Apoptosis | 18 | 0.000385 |
| Natural killer cell mediated cytotoxicity | 24 | 0.00139 |
| MAPK signaling pathway | 38 | 0.00297 |
| Leukocyte transendothelial migration | 19 | 0.00374 |
| RIG-I-like receptor signaling pathway | 11 | 0.0039 |

### Molecular pathogenesis of H5N1 infection in chicken

Previous transcriptomics study of our group found that highly pathogenic H5N1 virus induced excessive expression of type I IFNs, cytokine, chemokines and ISGs in the lung tissues (Ranaware et al., 2016). This atypical expression of immune genes (cytokine storm) might be the cause for the high mortality in chickens. However, the information on the pathways activated, constituents of cytokine storm and therapeutic strategies against cytokine storm is lacking in avian species. Intensive molecular studies in humans and human animal models system have identified, (1) Activation of TLR3 and 7, as well as endosome (TLR3 and 7) and cytosolic (RIG-I) pathways; (2) activation of IL-1R signaling pathway; and (3)activation of MVAS/MyD88/TRIF signaling as essential pathways involved in the cytokine storm (Teijaro et al., 2014).

In order to check whether these pathways are activated and if activated what are the levels of expression of the immune genes in chicken lung tissues, we applied meta**-**analysis of lung transcriptome datasets. We utilised the previously published microarray datasets of chicken infected with HPAIVs, because transcriptomic data can capture the complete gene expression dynamics at a particular condition. We mapped the differentially expressed genes retrieved from meta-analysis results of chicken into influenza reference pathways in KEGG database. Activation of TLR signaling pathway, RIG I signaling pathway, NOD like receptor signaling pathway and JAK-STAT signaling pathway were observed in chicken lung transcriptome (Figure S2). Similarly, activation of these pathways was evident in the chicken proteome datasets (Figure 3). Further Bayesian networks (BN) were constructed with prior knowledge using chicken meta-analysis transcriptome datasets (Figure 4, Figure S3). The combined *in silico* analysis of transcriptome and proteome datasets confirmed the activation of TLRs, RLRs, NLR and Jak-STAT signaling pathways in lung tissues infected with HPAIVs in chickens. The fact that most of the influenza pathogenesis possesses abnormalities in all of these core pathways suggests they play a central role in the cytokine storm.

**Fig 3.**
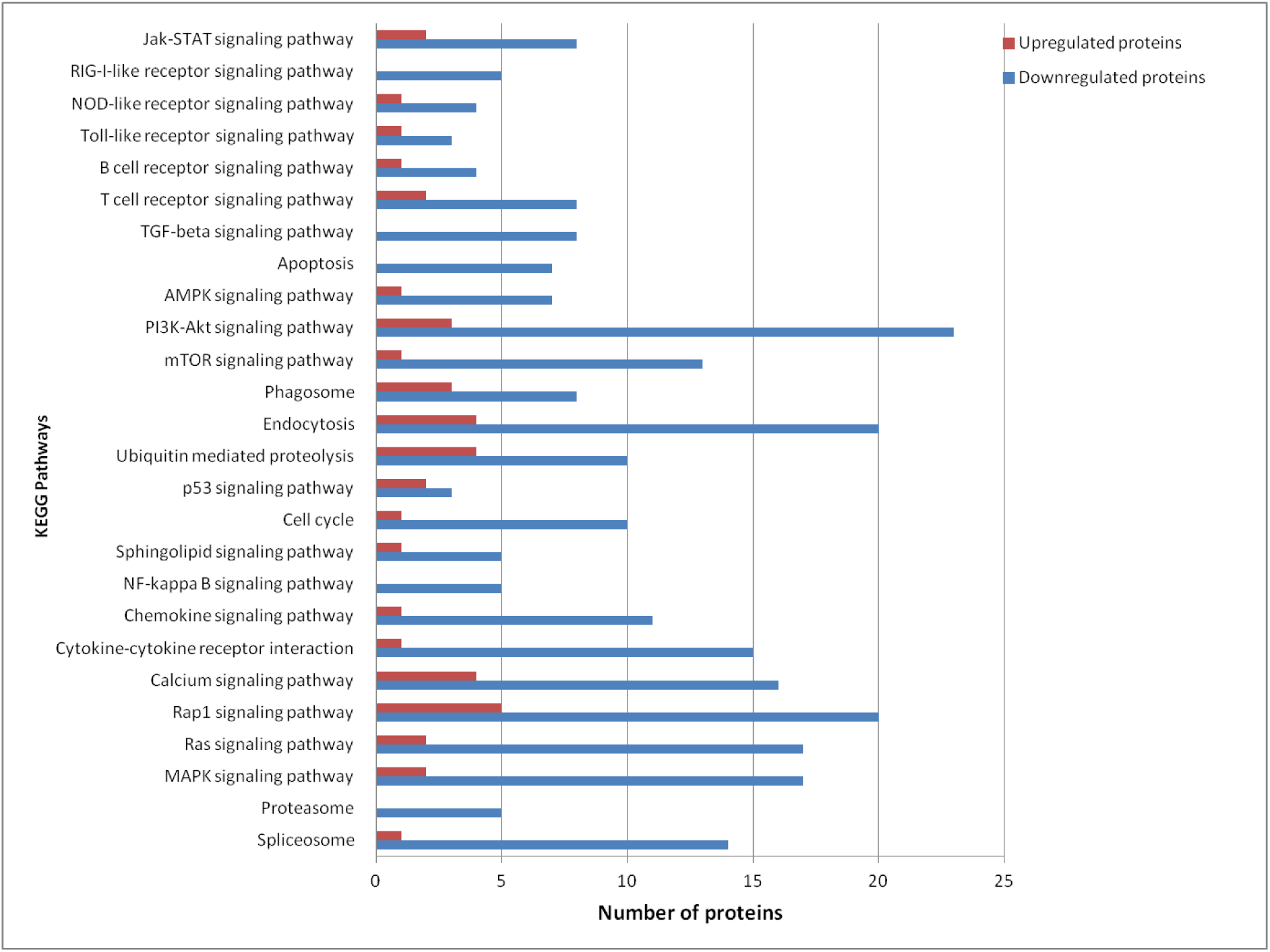
KEGG pathway analysis of commonly upregulated and downregulated proteins in chicken lung tissues infected with HPAI H5N1 virus.

**Fig 4.**
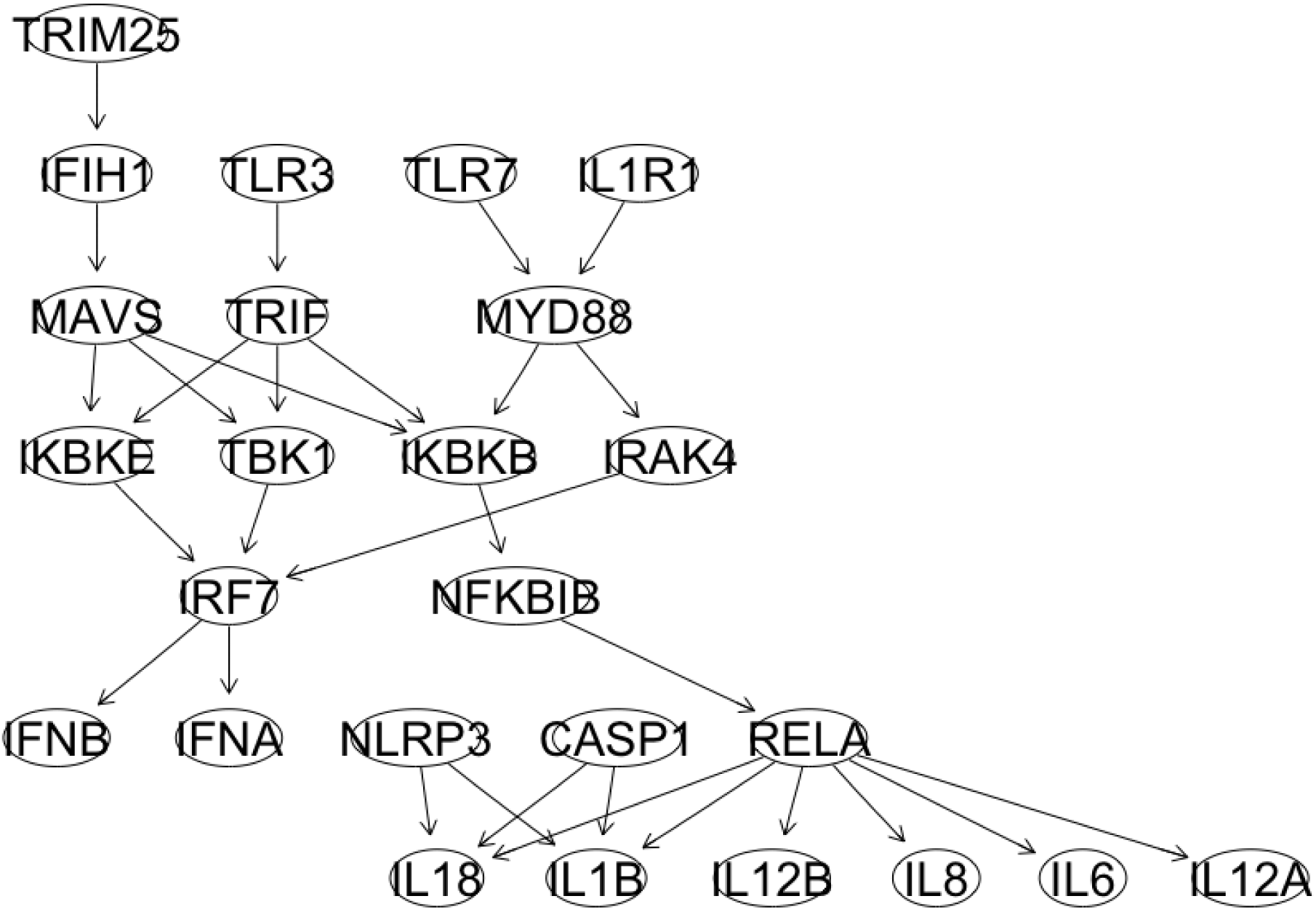
Bayesian network was constructed based on meta-analysis of transcriptome dataset of chicken. BN constructed with prior knowledge of TLR, RLR, IL1R and NLR signaling pathways in chicken.

Next we examined the cytokine storm responsive genes (i.e expression level of cytokines, chemokines and ISGs) as a result of activation of the above mentioned pathways. Cytokine storm responsive genes lists were compiled for chickens based on information in literature (Mishra et al., 2017). The expression levels (fold change) of these genes were retrieved from meta-analysis transcriptome datasets of chicken. Cytokines, chemokines and ISGs genes were found to be upregulated in chicken lung tissues and these may be the basis of the increase in the severity of HPAI H5N1 infection in chickens (Figure 5). In summary, we characterized the immune pathways involved in the cytokine storm and identified cytokine storm responsive genes in chicken lung tissues infected with HPAIVs.

**Fig 5.**
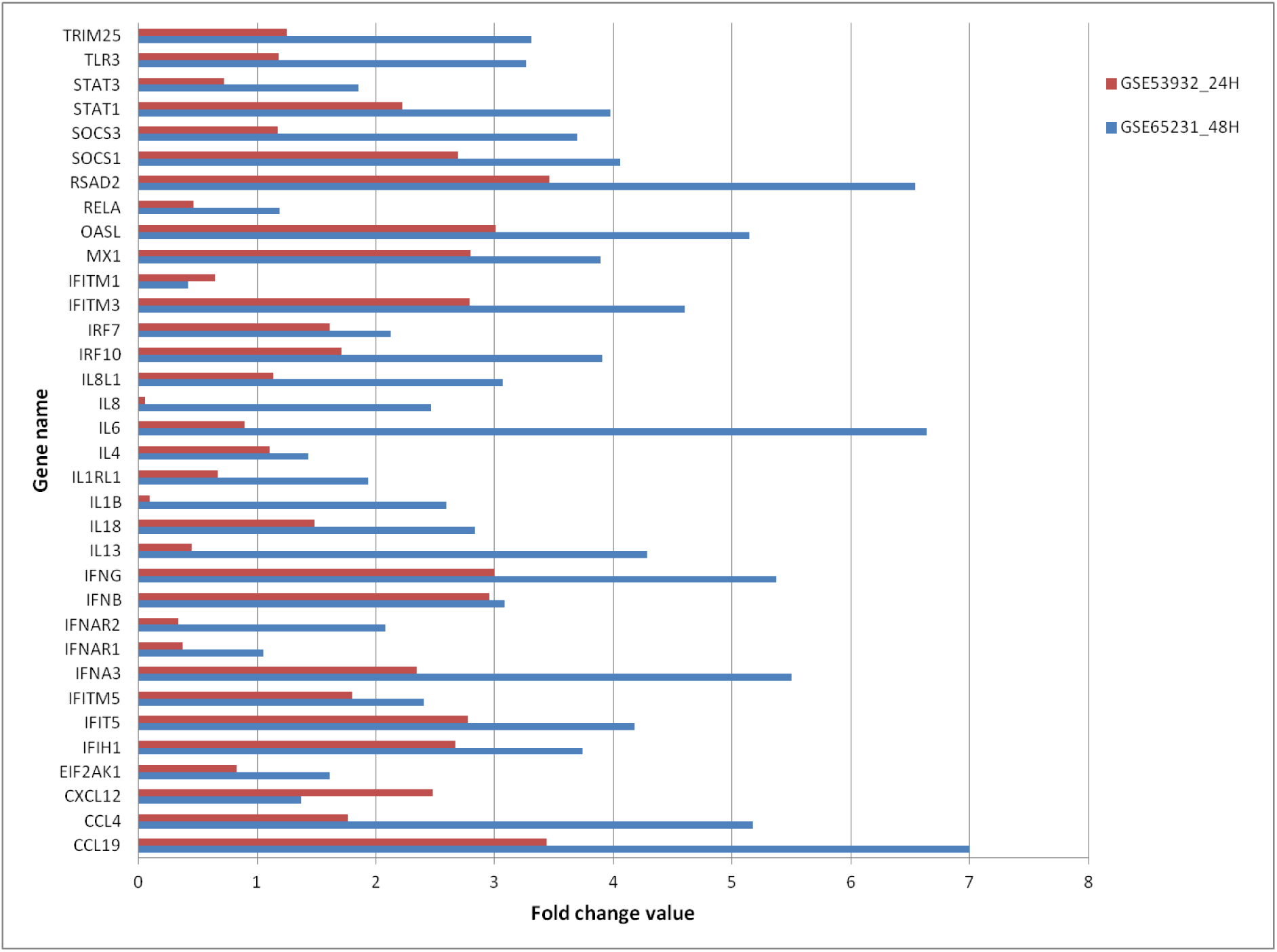
Expression level of cytokine storm responsive genes in chicken lung tissues infected with HPAIVs.

### Identification of proteomic determinants responsible for disease pathogenesis in chicken

To know the main driver or hub proteins responsible for disease pathogenesis, we constructed protein-protein interaction (PPI) network based on chicken lung proteome dataset (Figure 6). The proteins such as MyD88, IKBKB, IRAK4, RELA, and MAVS involved in the TLRs, RLRs, IL1R and NLR signaling pathways were identified with high degree centrality and betweenness centrality measures (Table 4).

**Table 4.** Hub proteins identified in chicken PPI networks based on degree centrality and betweenness centrality measures.

| **Proteins** | **Degree centrality** | **Betweenness centrality** |
| --- | --- | --- |
| HNF4A | 254 | 254342.21 |
| ELAVL1 | 254 | 253172.78 |
| FN1 | 117 | 79642.94 |
| COPS5 | 117 | 73208.97 |
| CUL1 | 105 | 55049.71 |
| CAND1 | 105 | 45936.7 |
| CTNNB1 | 80 | 75126.54 |
| BRCA1 | 72 | 41422.24 |
| FYN | 65 | 35789.72 |
| MYD88 | 10 | 2614.04 |
| IKBKB | 42 | 16555.52 |
| RELA | 48 | 23497.55 |
| MAVS | 6 | 1721.53 |
| STAT1 | 81 | 46280.33 |
| STAT2 | 14 | 1579.16 |
| STAT3 | 90 | 66853.77 |
| SOCS3 | 23 | 11524.2 |
| IRAK4 | 8 | 311.94 |

**Fig 6.**
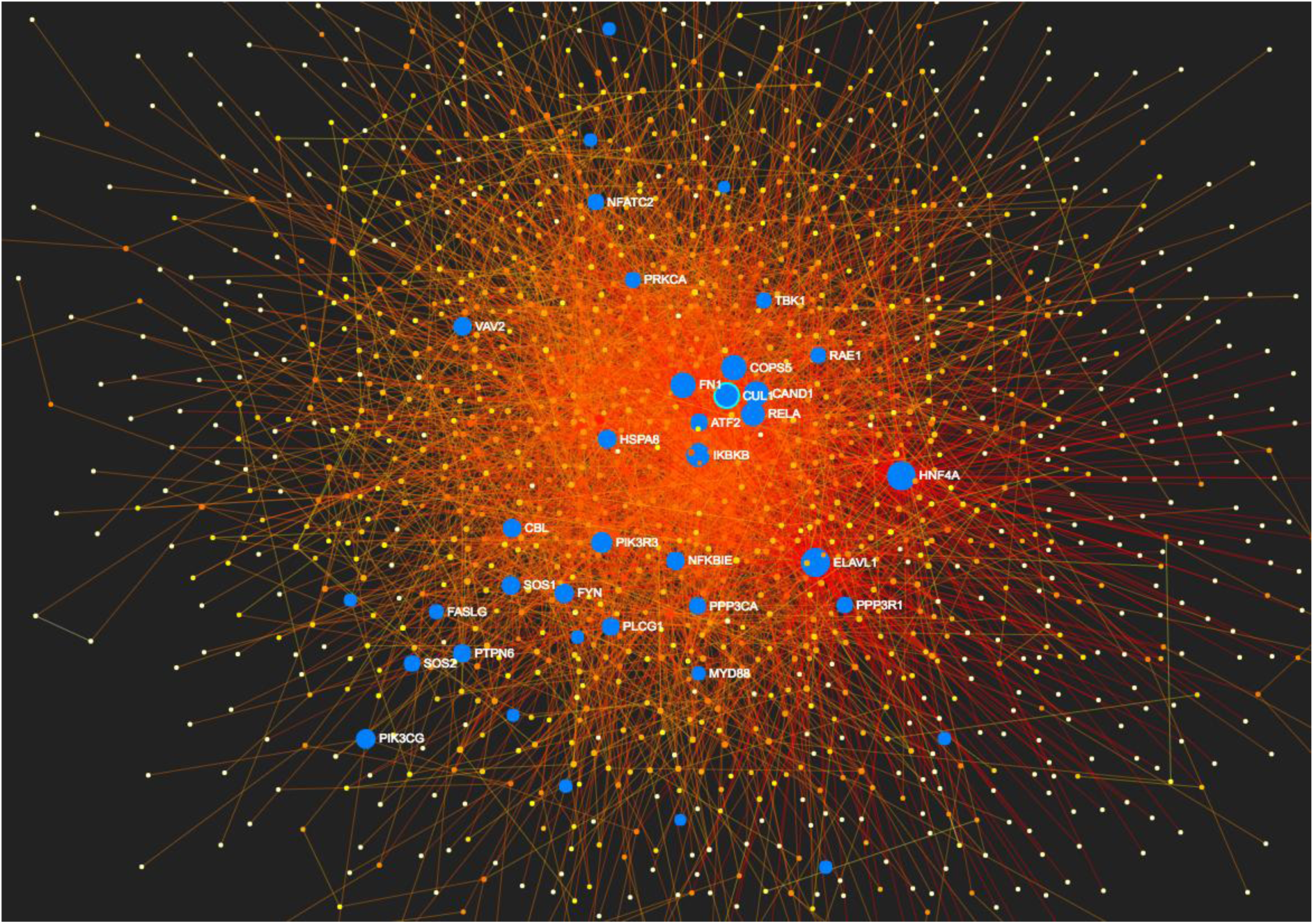
The protein-protein interaction network of chicken lung tissues infected with HPAI H5N1 virus. The important hub genes involved in influenza molecular pathogenesis were highlighted with blue colour in the PPI network.

MYD88 gene encodes a cytosolic adapter protein that plays a central role in the innate and adaptive immune response. This protein functions as an essential signal transducer in the IL1R and TLR signaling pathways. These pathways in turn regulate the activation of numerous proinflammatory genes (Deguine and Barton, 2014). The IKBKB protein phosphorylates the inhibitor in the inhibitor/NF-kappa-B complex, causing dissociation of the inhibitor and activation of NF-kB signaling pathway (Rothwarf and Karin, 1997). MAVS acts downstream of DDX58/RIG-I and IFIH1/MDA5 genes, as an essential signal transducer in the beta interferon signaling pathways and that contribute to antiviral immunity (Seth et al., 2005). RELA/NF-kB is a ubiquitous transcription factor involved in several biological processes. This transcription factor is activated through degradation of its specific inhibitor in the cytoplasm; NF-kB moves to the nucleus and activates transcription of specific genes. The NF-kBp65-p65 complex appears to be involved in invasin-mediated activation of IL-8 expression (Kunsch and Rosen, 1993). Teijaro et al. (2014) reported that MyD88 and MAVS as the predominant signaling molecule required for innate immune cell recruitment and for the majority of cytokine amplification (i.e. cytokine storm) in mice infected with influenza virus (Teijaro et al. 2014). Further they suggested that therapeutic control of cytokine storm is possible through a common pathway inhibition downstream of multiple innate pathogen-sensing molecules of cytokine amplification. In our study the identified hub proteins (MyD88, IKBKB, IRAK4, RELA, and MAVS) were all involved in different components of MyD88 and MVAS signaling pathways. Based on this literature, we suggest that successful therapeutic intervention for cytokine storm in chicken should target these proteins as drug targets to blunt the cytokine amplification. Further the S1P1R agonist therapy may suppress global cytokine amplification in chicken as in mice (Teijaro et al. 2014). However biological validation of this hypothesis by *in vivo* experiment is needed.

In Jak-STAT signalling pathway, we found STAT1, STAT2, STAT3 and SOCS3 proteins as the main driving proteins in the PPI network (Table 4). However, these proteins had protein probability range from 0.63 to 0.90, hence were not evident in our main proteome dataset of chicken. The STAT1, STAT2 and STAT-3 proteins are a key constituents of JAK-STAT signalling pathway, play critical roles in the IFN signalling pathway and are required for a robust IFN-induced antiviral response (Yang et al., 1998; Ho and Ivashkiv, 2006). SOCS1 and SOCS3 are reported to be critical regulators of IFN responses through the inhibition of STAT phosphorylation and induction of ISGs through a RIG-I/MAVS/IFNAR1-dependent pathway (Pauli et al., 2008; Pothlichet et al., 2008).

Further we identified some novel main driver/hub proteins with a very high degree of centrality and betweenness of centrality measures which are not reported previously to be associated with influenza infection condition in human, human models or avian species. These novel hub proteins are HNF4A, ELAVL1, FN1, COPS5, CUL1, CAND1, BRCA1, CTNNB1 and FYN (Table 4, Figure 6). The HNF4A protein is a transcriptionally controlled transcription factor and required for the transcription of alpha 1-antitrypsin, apolipoprotein CIII, transthyretin genes and HNF1-alpha genes (Ryffel, 2001). ELAVL1 is a member of the ELAVL family of RNA-binding proteins and selectively bind AU-rich elements (AREs) found in the 3’ untranslated regions of mRNAs. The ELAVL family of proteins play a role in stabilizing ARE-containing mRNAs (Brennan and Steitz, 2001). This gene has been implicated in a variety of biological processes and has been linked to a number of diseases, including cancer (Wang et al., 2013).

FN1 bind cell surfaces and various compounds including collagen, fibrin, heparin, DNA, and actin. FN1 is involved in cell adhesion and migration processes during embryogenesis, wound healing, blood coagulation, host defense, and metastasis (Liao et al., 2018). COPS5 is one of the eight subunits of COP9 signalosome and functions as an important regulator in phosphorylation of p53/TP53, c-jun/JUN, IkappaBalpha/NFKBIA, ITPK1 and IRF8 signalling (Bech-Otschir et al., 2002). CUL1 is a core component of multiple cullin-RING-based SCF (SKP1-CUL1-F-box protein) E3 ubiquitin-protein ligase complexes, which mediate the ubiquitination of proteins involved in cell cycle progression, signal transduction and transcription (Kipreos et al., 1996).

BRCA1 encodes a nuclear phosphoprotein that has a role in maintaining genomic stability, and acts as a tumour suppressor. This protein is involved in transcription, DNA repair of double-stranded breaks, and recombination (Foulkes and Shuen, 2013). FYN gene encodes a membrane-associated tyrosine kinase that has been implicated in the control of cell growth (Zheng et al., 2017). In summary, many proteins involved in the TLRs, RLRs, NLR and Jak-STAT signaling pathways and other novel proteins were identified as main protein determinant and these proteins might be linked with disease pathogenesis in chicken H5N1 infection. However critical functional role of these proteins in avian influenza pathogenesis in chicken requires further biological validation by *in vivo* and *in vitro* experiments.

In conclusion, we characterized the comprehensive proteome profile of chicken lung tissues infected with HPAI H5N1 virus at different time points post-infection. There are differences in the protein profile at different time points post-infection indicated by the remarkable variations in the levels of expression of proteins at these time points. Combined analysis of transcriptome and proteome datasets revealed activation of TLR, RLR, NLR, and JAK-STAT signaling pathways associated with a cytokine storm effect in chicken infected with HPAI H5N1 virus. Further, we identified many of the important hub proteins linked with influenza pathogenesis in chicken.

## Data availability

The mass spectrometry proteomics data have been deposited to the ProteomeXchange Consortium via the PRIDE partner repository with the dataset identifier PXD010358.

## Ethics statement

The experiments were approved by the Institutional Animal Ethics Committee of ICAR-NIHSAD (Approval no. 68/IAEC/HSADL/12 dated 11.05.2012), and performed under the guidance of the Committee for the Purpose of Control and Supervision of Experiments on Animals (CPCSEA), Ministry of Environment and Forests, Govt. of India.

## Acknowledgements

This work was funded by the Department of Biotechnology (grant number: BT/IN/Indo-UK/FADH/48/AM/2013). We thank Director, ICAR-National Institute of High Security Animal Diseases, Director, ICAR-Indian Veterinary Research Institute, and Indian Council of Agricultural Research, India for providing necessary facilities to carry out this work.

## Author Contributions

Conceived and designed the experiments: A.M., A.A.R. and P.V. Performed the experiments: A.M., S.C., A.A.R and P.V. Analyzed the data: P.V. and A.M. Contributed reagents/materials/analysis tools: H.V.M., D.D.K. and V.P.S Wrote the paper: P.V. and A.M.

## Conflict of Interest

The authors declare that the research was conducted in the absence of any commercial or financial relationships that could be construed as a potential conflict of interest.

