## Supplementary material for "Proteomics Analysis of Differential Expression of Lung Proteins in Response to Highly Pathogenic Avian Influenza Virus Infection in Chicken": Figure S1; Figure S2; Figure S3

\# Present affiliation of author: Veterinary College and Research Institute, Tamil Nadu Veterinary and Animal Sciences University, Orathanadu- 614625, Tamil Nadu, India

Figure S1. Representation of ProteinProphet predicted sensitivity and error rate. It is desirable for the red curve (sensitivity) to hug the upper right corner, and for the green curve (error) to hug the lower left corner.

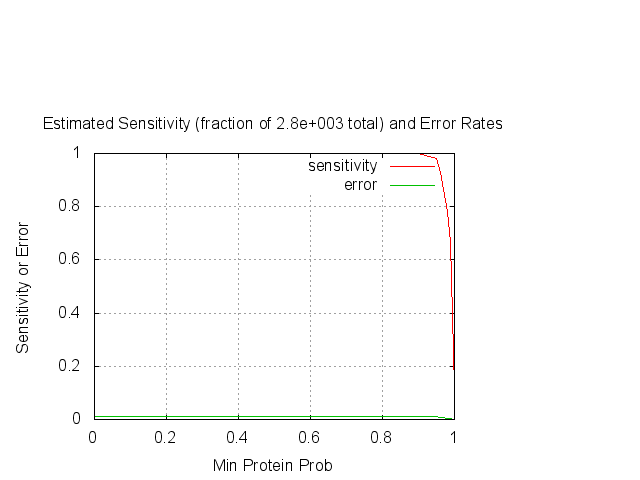

Figure S2. KEGG pathway mapping of differential expressed chicken genes list into influenza reference pathway in KEGG database^55^.

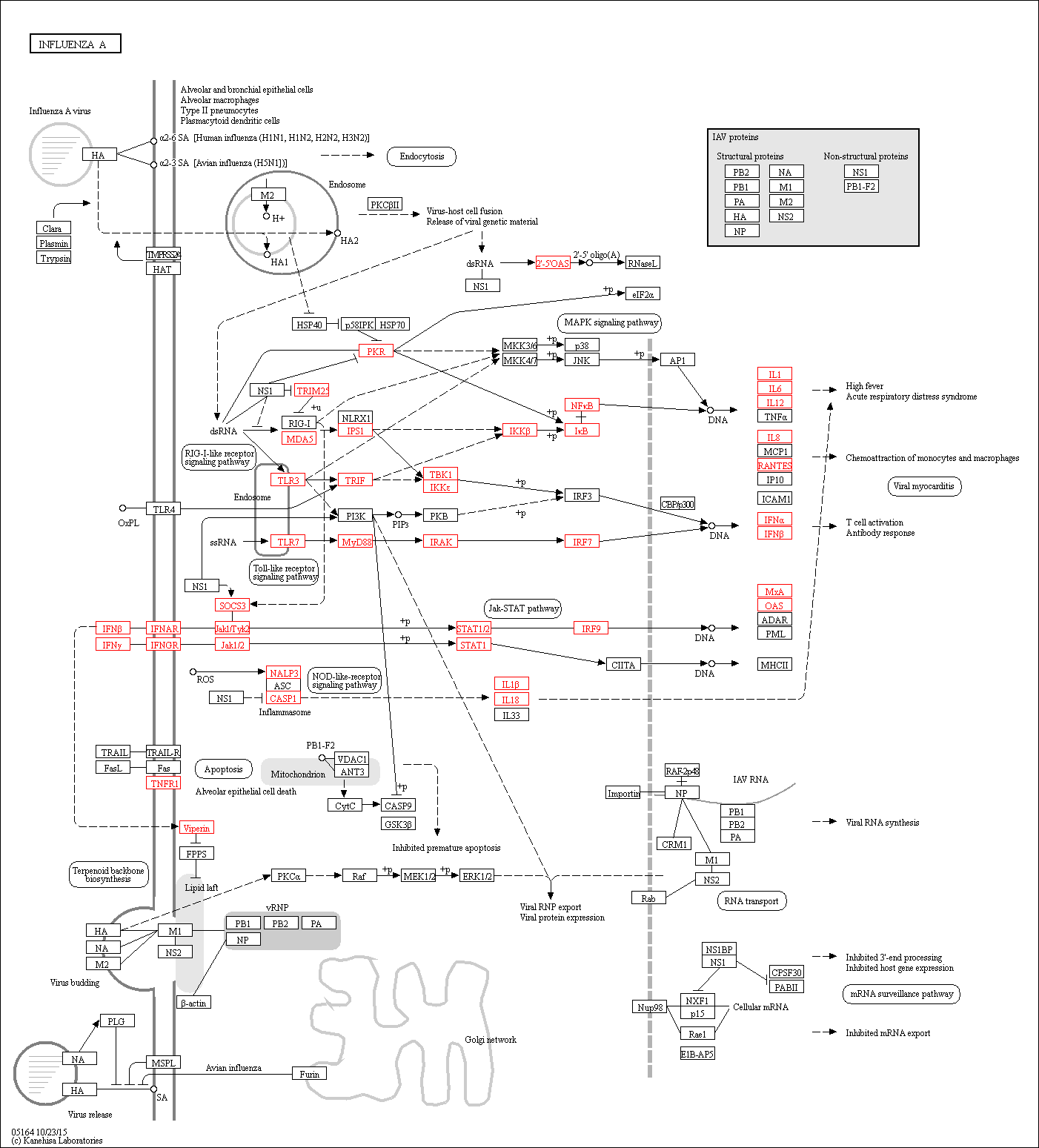

Figure S3. Bayesian network was constructed based on meta analysis transcriptome dataset of chicken for JAK-STAT signaling pathway with prior knowledge.

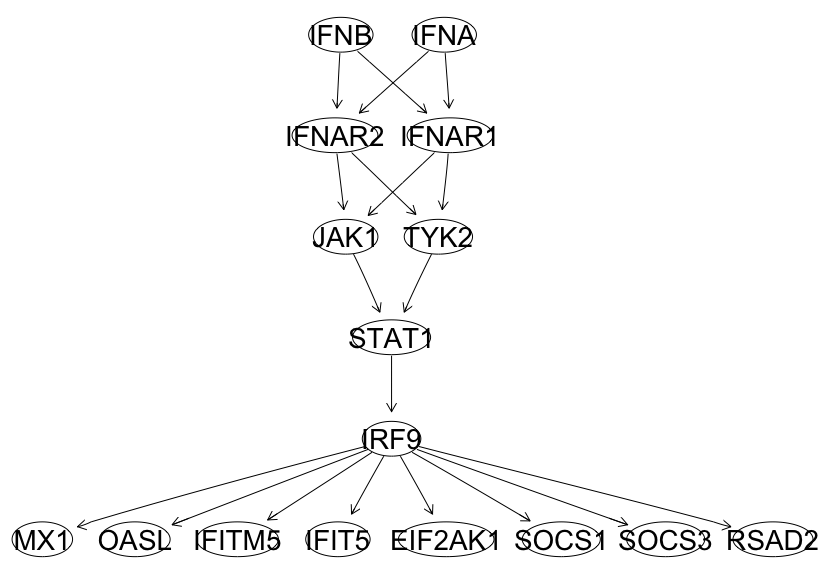
